# Interfacial dynamics and growth modes of *β*_2_-microglobulin dimers

**DOI:** 10.1101/2022.12.28.522115

**Authors:** Nuno F. B. Oliveira, Filipe E. P. Rodrigues, João N. M. Vitorino, Patrícia F. N. Faísca, Miguel Machuqueiro

## Abstract

Protein aggregation is a complex process that strongly depends on environmental conditions and has considerable structural heterogeneity, not only at the level of fibril structure but also at the level of molecular oligomerization. Since the first step in aggregation is the formation of a dimer, it is important to clarify how certain properties (e.g., stability or the interface geometry) of the latter may determine the outcome of aggregation. Here, we developed a simple model that represents the dimer’s interfacial region by two angles (spanning the so-called growth landscape), and investigate how modulations of the interfacial region occurring on the ns–*μ*s timescale change the dimer’s growth mode. We applied this methodology to 15 different dimer configurations of the *β*_2_m D76N mutant protein equilibrated with long MD simulations and identified which of them have limited and unlimited growth modes, with different consequences to their aggregation potential. We found that despite the highly dynamic nature of the starting configurations, most polymeric growth modes tend to be conserved within the studied time scale. The proposed methodology performs remarkably well taking into consideration that the *β*_2_m dimers are formed by monomers with detached termini, and their interfaces are stabilized by non-specific apolar interactions, leading to relatively weak binding affinities.

## Introduction

Protein aggregation is the process in which soluble protein conformations (monomers) with exposed hydrophobic patches, self-associate into dimers and higher-order oligomers.^1^ Aggregation may lead to amorphous aggregates with a granular appearance, protofibrils (including annular oligomeric aggregates), or other oligomeric aggregated states. Often, the end product of protein aggregation are amyloids, insoluble aggregates comprising long unbranched fibers, characterized by the cross-beta structure.^2^ While the formation of amyloids is the hallmark of a plethora of conformational disorders, including Parkinson’s and Alzheimer’s disease,^3^ it is thought that the most cytotoxic species within the aggregation cascade are the early-forming oligomers. Indeed, the latter not only cause an overload of the proteostasis machinery, but can also seed the aggregation of identical proteins, cross-seed the aggregation of homologous proteins, and even disrupt biological membranes (reviewed in^4^).

Since protein aggregation typically starts with the self-association of two monomers into a dimer, by blocking dimerization one could - at least in principle - hinder aggregation. Therefore, it is critical to have a molecular-level understanding of the mechanism of dimer formation and to establish how certain traits of the dimer (including structure, conformational dynamics, energetic and thermodynamic stability, etc) may determine the outcome of aggregation. It has been suggested that a properly formed dimer with increased stability may be protective against aggregation.^5^ Additionally, it has been observed that a larger conformational variability at the dimer level appears in association with a faster aggregation process and higher toxicity.^6^

The identification and structural characterization of early-formed oligomers is particularly challenging because they exist in a complex dynamic equilibrium with each other and with insoluble higher order aggregates.^7,8^ An additional difficulty is that they form transiently, in a timescale that is not compatible with the temporal resolution of commonly used biophysics apparatus. Furthermore, their formation is under kinetic control, being strongly dependent on environmental conditions. ^9,10^ Computational modelling covering different resolutions and timescales has been playing a pivotal role in helping understand the nature of protein aggregation^11^ and in providing predictions for the structure of the oligomeric species formed along the amyloid pathway.^7,12,13^

In recent years, some of us developed an integrated computational methodology (reviewed in^14^) combining data from discrete molecular dynamics (DMD) of a structure-based model,^15–17^ constant-pH MD,^18–21^ and a rigid-body Monte Carlo ensemble docking method developed in house (MC-ED)^15^ that uses a low-resolution cost function based on step potentials to model electrostatic, hydrophobic and hydrogen bond interactions.^22^ The MC-ED provides a probabilistic view of the dimerization phase, in particular, which regions of the monomer are more likely to associate, and which residues play a key role in aggregation acting as aggregation hot spots. We applied it to study the dimerization phase of protein beta-2-microglobulin (*β*_2_m) under different environmental conditions,^14^ but the method is general and, in principle, can be used for any model system.

*β*_2_m is a small (99 residues long) protein with biomedical interest.^14^ It has a classical beta-sandwich fold in which the native structure is stabilized by a disulfide bridge between the sulfur atoms of the cysteine residues 25 (at B strand) and 80 (at F strand)^23^ (Figure 1A). The disulfide bridge has been considered critical for *β*_2_m fibrillogenesis.^24,25^ The wild-type form (wt-*β*_2_m) is the canonical agent of dialysis-related amyloidosis, a disease that affects individuals with kidney impairment undergoing long-term hemodialysis,^26^ while the single point mutant D76N is the causing agent of familial amyloidosis affecting the visceral organs.^27^ By exploring the folding transition of D76N with DMD simulations of a structure-based Gō model,^28^ we were able to predict an intermediate state with exposed hydrophobic patches (i.e., with the potential to trigger self-association), which is structurally characterized for having a well-preserved core and two unstructured and detached terminal regions.^28^ For this reason we termed this intermediate I_2_ (Figure 1B). Afterwards, a study that combined *in crystallo* with *in silico* analyses,^29^ reported a highly dynamic conformation, for D76N where the loss of beta structure at both the N- and C-terminal strands causes the exposition of very aggregation-prone regions, in consonance with our results. More recent DMD folding simulations indicate that the wt-*β*_2_m can also populate a similar conformer provided the disulfide bond is established.^30^ While the role of the experimentally detected intermediate state in the aggregation pathway of D76N still lacks consensus, ^32^ we decided to use the topologically similar I_2_ as a model system to explore the dimerization phase of *β*_2_m.^14^ By conducting extensive protein-protein docking simulations, we found a prominent role for the DE- and EF-loops as adhesion regions, with the N-terminal region becoming also relevant at acidic pH.^28^ Our analyses also predict that Trp60, located at the apex of the DE-loop, is no longer the leading hot-spot at acidic pH, losing its role for Arg3 located at the N-terminus.^22^ Given the coarse-grained nature of the deployed cost function, the MC-ED method should not be used to predict accurate structural dimer models. Motivated by this limitation, we recently developed a computational protocol that combines relatively short (100 ns) Molecular Dynamics simulations with MM-PBSA calculations. ^13^ The MD simulations correct structural errors (e.g., steric clashes), while the MM-PBSA method provides accurate interfacial energies for the relaxed dimers. Additionally, by using PyMol^33^ and a simple geometric protocol to replicate the dimer’s interface, we could determine by visual inspection which binding modes (i.e., dimer interfaces) lead to self-limited growth (i.e., polymeric chains that close upon themselves), and unlimited growth (i.e., polymeric chains that can growth indefinitely) in the absence of structural rearrangement. By deploying this methodology to an ensemble of 212 dimers obtained with the MC-ED, we found that the dimers are mainly stabilized by apolar interactions, and that both the N- and C-terminus are involved in the interface of the 10 most stable dimer interfaces.^13^ We also found that some of the considered dimer interfaces, including the most energetically stable, lead to self-limited growth, while other can growth indefinitely in the absence of structural rearrangement.

**Figure 1:**
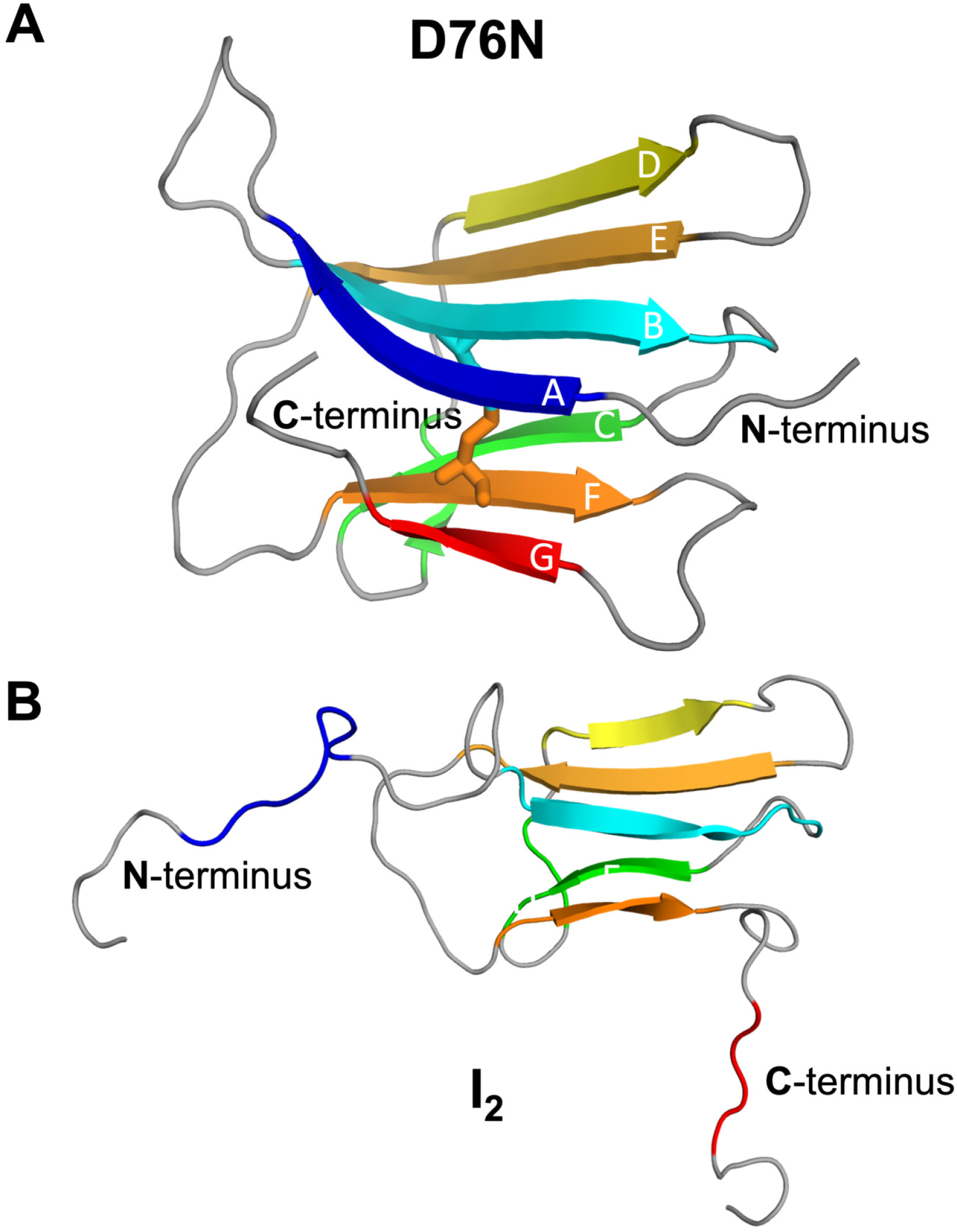
Three-dimensional cartoon representation of the native structure of D76N (pdb id: 2YXF^31^)(A) and the *I*_2_ intermediate state populated by D76N, which features a well-preserved core and two unstructured and decoupled termini (B).^28^

Here, we extend our previous study^13^ of *β*_2_m self-association and investigate how the growth mode is modulated by the geometric changes occurring at the dimer’s interface in the ns–*μ*s timescale. To achieve this, we developed a method that determines the growth mode based on just two angles, which constitute a minimal representation the dimer’s interfacial geometry. We consider the ten most stable binding modes identified in,^13^ as well as five additional binding modes of intermediate and lower stability. We observe that for most binding modes and despite significant changes in the geometry of the interfacial region, the limited/unlimited nature of the growth mode is conserved within the considered timescale.

## Methods

### Molecular Dynamics simulations

All molecular dynamics (MD) simulations were performed using GROMACS 2018.6,^34^ the GROMOS 54A7 force field,^35,36^ and the SPC water model.^37^ The starting conformation of our simulations is a relaxed homodimer of the I_2_ intermediate, ∼20k water molecules, and 2 Na^+^ ions. We chose a set of 15 dimers previously investigated in^13^ that consisted of: the top 10 dimers with the lowest binding energies (best affinity) (∼ −83 to ∼ −67 kcal mol^−1^); 2 dimers with very high energies (weak affinities; ∼ −10 kcal mol^−1^); and 3 dimers with average binding energies (∼ −43 kcal mol^−1^). To improve our sampling, we performed 3 replicates of 500 ns per dimer. Starting from the dimer simulations previously published,^13^ the first replicate went straight into production. The starting structures from the second and third replicates were obtained by performing a new velocity initialization (50 ps with a random seed) before proceeding to the long production runs. The production MD was conducted with a 2 fs time step and the electrostatics were treated with the Particle-Mesh Ewald (PME) method,^38,39^ with a verlet scheme cutoff of 1.4 nm, a Fourier grid spacing of 0.12 nm, and an interpolation order of 4 (cubic). Van der Waals interactions were simply truncated above 1.4 nm.^40^ The neighbor list was updated every 10 integrator steps. The LINCS algorithm^41^ was used to constrain all the protein bonds and SETTLE was used on the water molecules.^42^

### Structural analysis

All structural analyses of the dimers were carried out using the GROMACS package and other in-house tools. The periodic boundary conditions effects were corrected using the fixbox tool.^43^ When dealing with equilibrium properties, the first 250 ns of the simulation were discarded. For the structural alignments and Root Mean Square Deviation (RMSD) calculations, only the Cα atoms from the protein core were considered (namely, residues 23-27, 36-39, 51-55, 62-66, and 78-82 of each monomer). This approach discards spurious structural variability due to loops and unstructured N- and C-terminal regions. As the RMSD reference structure, we used the initial structure, which corresponded to the relaxed structure obtained in our previous work.^13^

The interfacial area of the dimers was calculated using the difference between the SASA values of both monomers in the presence and absence of the partner.^44,45^ This procedure was also applied to each residue individually, with the final area being normalized (converted into a percentage) by the maximum SAS area of each residue type observed in all simulations. Residues with interfacial area higher than 10%, were considered to be located at the interface.

The asphericity of each monomer was calculated using the radius of gyration (*R*_*g*_) and the eigenvalues of the gyration tensor (λ_*i*_), given by the square of the three principal radii of gyration (*R*_*x*_, *R*_*y*_, *R*_*z*_) obtained using the GROMACS tool *polystat*. The asphericity of each monomer, *A*_*s*_, is given by the equation 1. ^46,47^

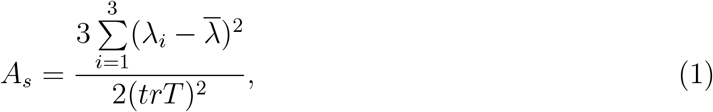

where 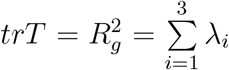 and 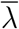 is the average eigenvalue of the inertia tensor given by 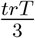Values of A_*s*_ higher than 0 indicate a deviation from the ideal sphere shape of our protein. Since 0 ≤ A_*s*_ ≤ 1, we can obtain the sphericity, ψ, in percentage, by taking (1 − *A*_*s*_) × 100. To obtain the average structures of each equilibrated dimer replicate, the GROMACS clustering tool was employed with an RMSD cut-off of 2 nm, following the GROMOS clustering algorithm.^48^

All error bars indicate the standard error of the mean. Images were rendered using PyMOL^33^ and Blender.^49^

### Calculation of binding energies with PyBindE

All binding free energy (E_*bind*_) calculations were performed using PyBindE, a new in-house implementation of the Molecular Mechanics Poisson-Boltzmann Surface Area (MM/PBSA) method,^50,51^ written in the programming language Python (https://github.com/mms-fcul/pybinde). We used a dielectric constant of 4, a grid scaling factor (*scale*) of 2, and a convergence criterion (*convergence*) of 0.01 kT/e.^13^ Additionally, the Poisson–Boltzmann (PB) calculations used 500 and 50 linear (*nlit*) and non-linear (*nonit*) iterations, respectively. The grid size was calculated as double the maximum value of the 3 atomic spatial coordinates, plus 1 if the final number is even, while the center was located in the geometric center of all the atomic coordinates of the system. For each dimer, the MM-PBSA calculations were performed every 100 ps, which resulted in 3× 5000 frames per dimer.

### A simple protocol for predicting and visualizing dimer replication

In our previous study^13^ we deployed a simple replication protocol that allows predicting the three-dimensional structure of a polymerized chain of *β*_2_m dimers forming in the absence of structural rearrangement. In particular, we used PyMOL to visualize the structure of a polymerised chain that was created by the repeated addition of dimers following a simple geometric protocol (Fig 2). In doing so, we found that some dimer interfaces lead to chains that close upon themselves (self-limited growth), while other interfaces can sustain indefinite growth.^13^

**Figure 2:**
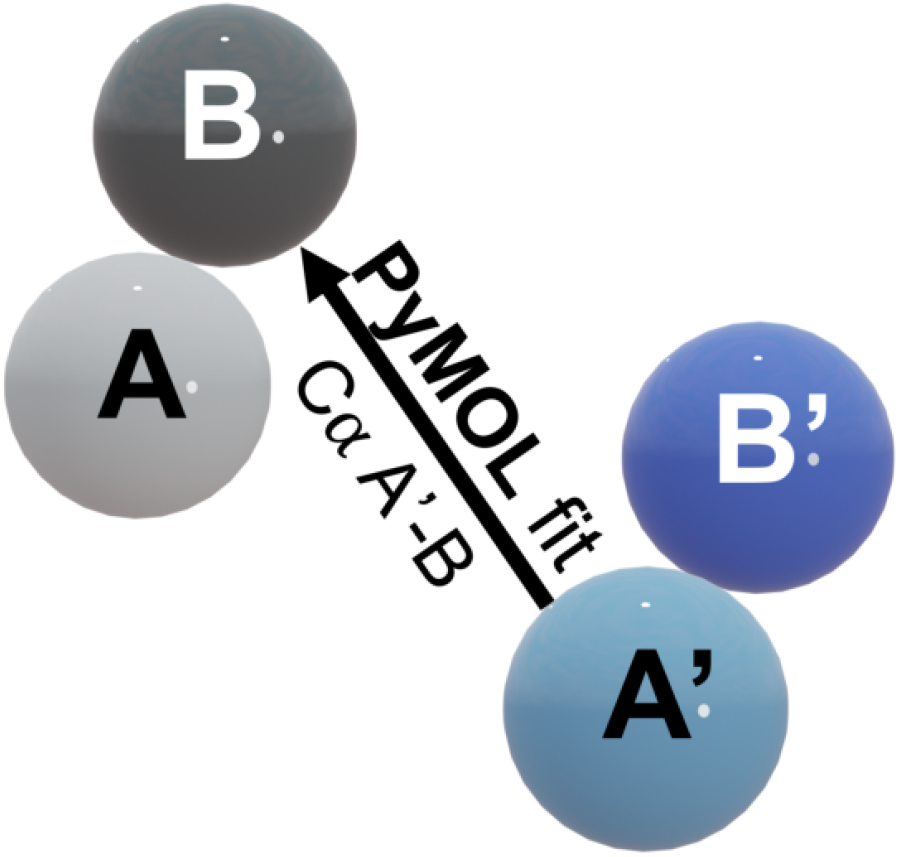
A simple protocol for dimer duplication and polymeric growth. To generate a polymerised chain of dimers, we consider a starting dimer conformation formed by monomers A and B. We subsequently create a duplicate dimer conformation where monomers A and B are respectively renamed A′ and B′. We then use PyMOL to structurally align monomer A′ from the duplicate dimer with monomer B of the preceding dimer and repeat this operation a desired number of times.

### A simple model for dimer polymerization

In the present study, we develop a simple model for chain polymerization and combine it with the protocol for dimer replication. In doing so, we are able to predict in a systematic manner which dimer interfaces lead to self-limited or indefinite growth modes. The model represents the dimer by two spheres of unit radius (one sphere per monomer) and reduces the description of dimer polymerization to two angles: a polymerization angle (*θ*_*pol*_), which determines the structure of the polymerized chain, and a polymerization dihedral angle (*ϕ*_*pol*_), which establishes the relative orientation of the two monomers in the dimer. The polymerization angle is defined by

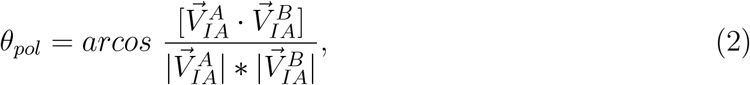

where 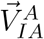 is the vector between the geometric center of monomer A (GC_*A*_) and the geometric center of the set of interfacial residues of A which are in contact with monomer B (GC_*IA*_), and 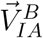 is the vector between the geometric center of monomer B (GC_*B*_) and GC_*IA*_ mapped onto the surface of monomer B (Fig 3A,B). On the other hand, the polymerization dihedral is given by

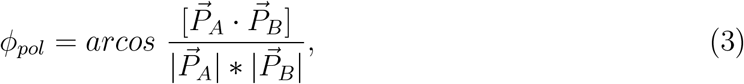

where 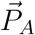 is the normal vector of the plane defined by the vector between GC_*A*_ and GC_*B*_ 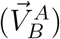, and the vector between GC_*A*_ and GC_*IB*_ mapped on monomer A 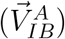. Similarly, 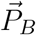 is the normal vector of the plane defined by GC_*B*_ and GC_*A*_ 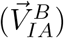 and the vector between GC_*B*_ and GC_*IA*_ mapped onto monomer B 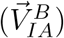 (Fig 3C).

**Figure 3:**
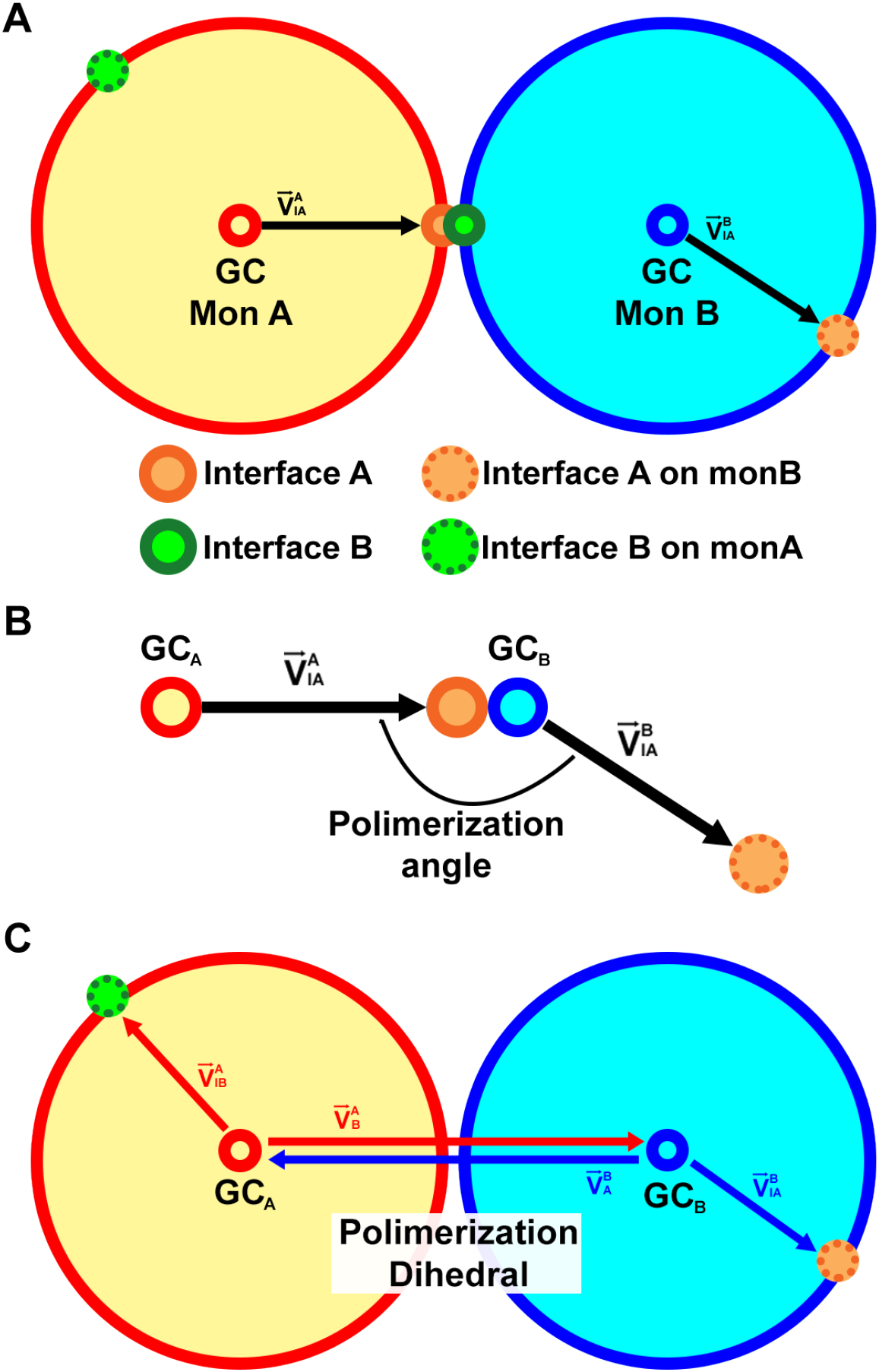
Schematics of points and vectors (A) used in the calculation of the polymerization angle, *θ*_*pol*_ (B), and dihedral, *ϕ*_*pol*_ (C). Monomer A and its geometric center are respectively represented by a large and small yellow sphere with a red contour, while monomer B and its geometric center are respectively represented by a large and small cyan sphere with a blue contour. The geometric center of the contact interfaces of monomers A and B are represented by small spheres with orange and green contours, respectively, while the projection of their interfaces on the opposing monomer are represented as small spheres with a dotted contour colored orange and green, respectively. On the polymerization dihedral red arrows represent the vectors used to calculate the normal vector 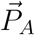 and the blue arrows represent the vectors used to calculate the normal vector 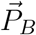.

Consider two initial spheres representing two monomers A and B. One of the spheres, say monomer A, has its center placed at the origin of a Cartesian reference frame, while the other has its center placed at (−2,0,0). A pair of angles (*θ*_*pol*_, *ϕ*_*pol*_) is chosen for this dimer. By duplicating the dimer thus created, one can use the previously described geometric protocol that aligns sphere A′ from the duplicate dimer with monomer B of the preceding one (Fig 2). By iterating this procedure a certain number of times one obtains a sphere model for dimer growth (Figure 4). By using this simplified model and the protocol for dimer replication, it becomes possible to map out self-limited and indefinite growth modes for a pair of parameters (*θ*_*pol*_, *ϕ*_*pol*_) (Figure 5). Considering several (*θ*_*pol*_, *ϕ*_*pol*_) pairs separated by 5° within the range 0°–180°, we were able to generate several models of a polymerized chain and identify its growth mode. A growth mode is classified as limited for the models exhibiting overlapping spheres. Within the 0°–60° range, the polymerization mode is, in general, of limited type, while the combination of *θ*_*pol*_ >60° with some dihedral values leads to unlimited growth. Based on this analysis, we created what we term by *growth landscape*, a two-dimensional surface generated by *θ*_*pol*_ and *ϕ*_*pol*_ that outlines the regions where growth is limited, uncertain, or unlimited (Figure 5). The transition points between limited and unlimited growth modes are shown as dots, and we used an exponential fit to this data set to draw the delimitation region.

**Figure 4:**
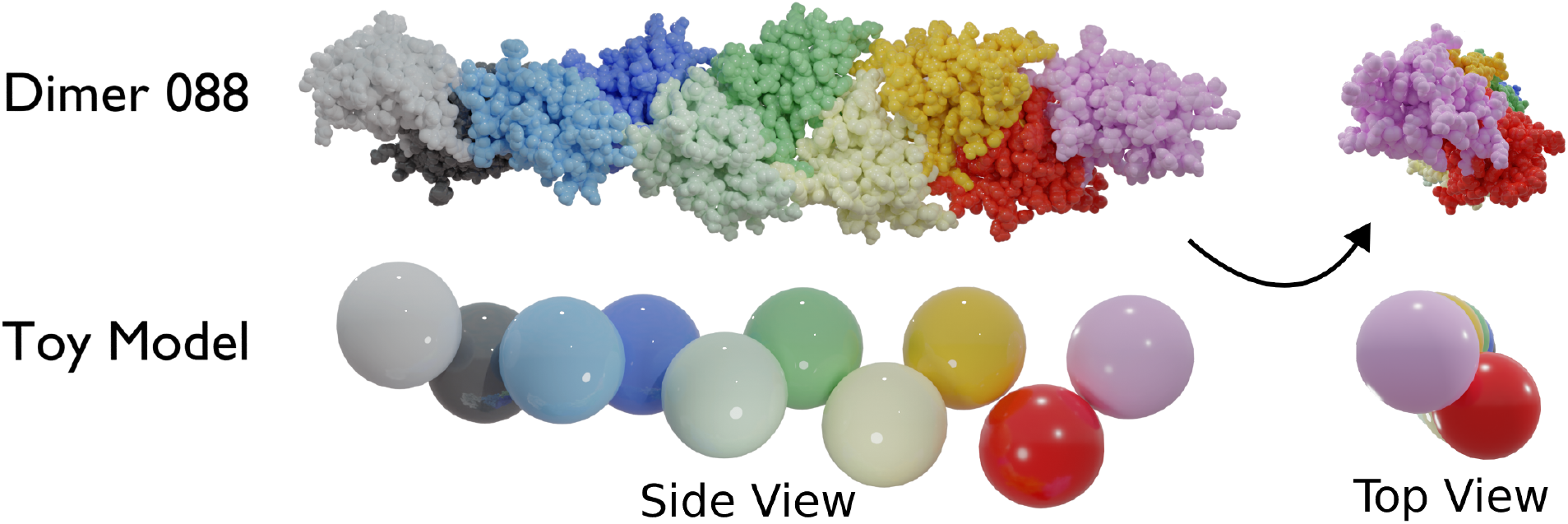
Structural polymerization of a selected dimer interface with *θ*_*pol*_=81° and *ϕ*_*pol*_=160° (BM-3, replicate 2) and corresponding sphere model. The original monomer A and monomer B are represented in white and black, respectively, and the consecutively added monomers follow the colors light-blue, blue, light-green, green, light-orange, orange, red, and pink. A representation of the side and top view of the monomer is given side by side.

**Figure 5:**
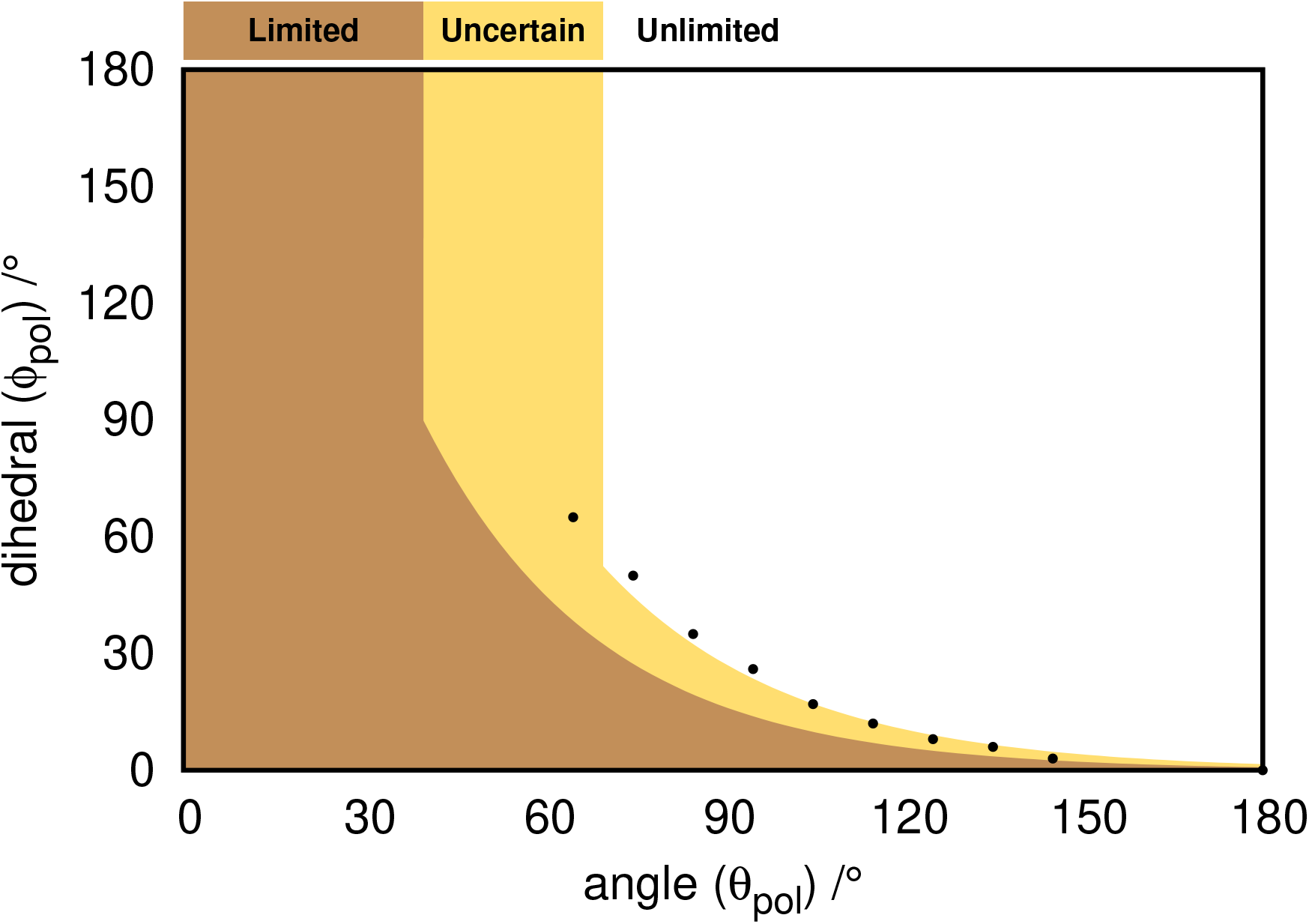
Growth landscape as a function of *θ*_*pol*_ angle and *ϕ*_*pol*_ dihedral angle. The black dots represent the upper limit of angle/dihedral combinations that yield a limited growth polymer as predicted by the simple model. The orange region represents the region where polymer growth is limited, the yellow color represents the uncertain region (one that requires visual inspection), and the region in white represents the unlimited growth.

This simple approach allows one to predict the type of growth mode associated with a *β*_2_m dimer conformation sampled in the course of a MD simulation because it suffices to calculate the angles *θ*_*pol*_ and *ϕ*_*pol*_ for that particular conformation. When a dimer falls into the uncertain region, it is necessary to visually confirm the type of growth mode by using the previously described protocol for dimer replication. We note that the observed uncertainty is mainly due to deviations of the monomer from the spherical shape considered in the simple model. Those deviations can be quantified by the monomer’s sphericity described in the previous section.

## Results and Discussion

### Binding energy and structural analysis

In our previous work,^13^ we performed 100 ns MD simulations to relax an ensemble of 212 *β*_2_m dimer configurations obtained with the MC-ED method (the *Relax* configurations). We evaluated the dimer’s binding energies, investigated which regions of the monomer are most likely present in the dimer’s interface, identified the so-called hot-spot residues (the residues that establish a higher number of intermolecular interactions), and determined which (of the most stable) interfaces could be propagated indefinitely allowing the formation of long, polymerized chains. Here, we will study *long-term* dimer stability and determine to which extent the latter is a determinant of the polymerization growth mode. This is an important question, whose answer will contribute towards a better understanding of heterogeneity in protein aggregation. In doing so, we will also address the methodological problem of gauging the quality of dimer conformation obtained with the MC-ED, which uses a low-resolution cost function to generate a dimer interface. Indeed, if a short MD simulation proves to be enough to relax the dimer conformation obtained with MC-ED, and establish its polymerization growth mode, the dimer conformation can be considered as a good predictive model. This is of practical importance since the MC-ED provides a dimer interface at a low computational cost.

To explore these questions, we performed long (3×500 ns) MD simulations of a subset of 15 dimers (the *Equil* conformations) that includes the 10 dimers with the lowest binding energy, 3 dimers with average binding energy, and the 2 dimers with highest binding energy from the ensemble studied in ^13^ (Table 1). We generically refer to these dimers as binding modes (BMs). For each dimer, we evaluated the binding energies together with several structural properties, such as RMSD, interfacial area, and secondary structure content. Perhaps not surprisingly, the most stable BMs equilibrate relatively fast (100 ns), with the MD simulations starting from the most unstable BMs requiring more time to equilibrate (Figures S1-S4 of Supporting Information). The RMSD (Figure S1 of Supporting Information) is highly sensitive to small structural and conformational rearrangements, making it challenging to achieve equilibration and/or replicate convergence. Nevertheless, even the most dynamic BMs/replicates roughly equilibrated within ∼250 ns as hinted even by their RMSD time evolution. Furthermore, to quantify the structural stability of the interfacial region in the long MD simulations, we calculated the percentage of residues, present in the relaxed interface,^13^ that are conserved after the 500 ns equilibration simulation (Table 1).

**Table 1.** Properties of *β*_2_m dimers. Binding energies, interfacial area and percentage of conserved interface for the set of 15 investigated dimers. The *Relax* data was obtained from our previous work,^13^ while the *Equil* data refers to the equilibrated segment (last 250 ns) of the long MD simulations carried out in the present study. The % shown in parenthesis in the *Equil E*_*bind*_ corresponds to the variation from the *Relax* value.

| BM | $E_{bind}$ (*Relax*)<br>(*kcal/mol*) | $E_{bind}$ (*Equil*)<br>(*kcal/mol*) | *Equil* Interf.<br>Area (*nm*<sup>2</sup>) | Conserved<br>Interf. (%) |
| --- | --- | --- | --- | --- |
| 1 | -82.9 ± 7.1 | -91.9 ± 7.3 (−11%) | 15.2 ± 1.0 | 76.4 ± 0.4 |
| 2 | -82.1 ± 5.2 | -100.8 ± 4.3 (−23%) | 17.5 ± 3.3 | 72.1 ± 3.7 |
| 3 | -79.0 ± 3.2 | -70.9 ± 1.9 (+10%) | 10.7 ± 0.6 | 70.5 ± 3.3 |
| 4 | -77.3 ± 7.6 | -86.2 ± 2.0 (−12%) | 13.3 ± 0.7 | 81.2 ± 2.1 |
| 5 | -75.4 ± 1.3 | -87.3 ± 5.1 (−16%) | 12.3 ± 2.8 | 85.4 ± 3.6 |
| 6 | -71.8 ± 13.4 | -74.9 ± 4.4 (−4%) | 13.0 ± 1.0 | 72.7 ± 1.3 |
| 7 | -71.0 ± 6.9 | -80.2 ± 8.9 (−13%) | 15.3 ± 3.0 | 75.1 ± 5.0 |
| 8 | -68.0 ± 3.3 | -55.5 ± 3.0 (+18%) | 11.9 ± 1.1 | 73.2 ± 1.4 |
| 9 | -67.5 ± 3.2 | -76.8 ± 7.1 (−14%) | 12.0 ± 1.5 | 61.2 ± 6.5 |
| 10 | -67.2 ± 2.0 | -70.2 ± 7.2 (−5%) | 12.7 ± 2.1 | 79.6 ± 2.6 |
| A-1 | -45.5 ± 0.6 | -55.1 ± 2.1 (−21%) | 10.3 ± 2.2 | 84.1 ± 5.3 |
| A-2 | -43.7 ± 1.8 | -53.1 ± 2.3 (−22%) | 10.3 ± 1.4 | 85.9 ± 4.1 |
| A-3 | -43.4 ± 2.3 | -59.4 ± 8.5 (−37%) | 10.2 ± 2.5 | 65.7 ± 2.9 |
| H-1 | -12.8 ± 1.7 | -14.1 ± 1.8 (−10%) | 6.3 ± 0.4 | 69.5 ± 4.8 |
| H-2 | -10.8 ± 1.3 | -16.7 ± 7.0 (−55%) | 5.8 ± 0.3 | 66.4 ± 7.8 |

With the exception of BM-3 and BM-8, whose binding energies increased by 10% and 18%, respectively, after the long MD equilibration, all the other dimer interfaces considered in this study became more stable, with the highest increase in stability (55%) being observed for H-2, followed by A-3 (37%) and BM-2 (23%). The energetic stability of the interfaces is related to their size, with the most stable BMs typically exhibiting the highest interfacial area. This is consistent with our previous finding that the van der Waals interactions are a major source of stability for these dimer interfaces.^13^ With the exception of BM-9, all the other most stable BMs were able to conserve a moderate (70%) to high (85%) fraction of their starting interface, which can thus be classified as being moderately stable from a structural point of view. Interestingly, although the two less stable dimers (H-1 and H-2) were able to decrease their binding energies after the full equilibration, the 500 ns timescale was not long enough to observe the structural rearrangement required to significantly decrease their binding energy to values comparable with BMs 1-10.

To gauge the quality of the data obtained from the previous MD relaxation step,^13^ we calculated the correlation between the binding energies and the average binding energies during the long equilibration step (calculated in intervals of 10 ns) (Figure 6). The high correlation values observed even after 500 ns of MD simulations indicate that, although the binding interfaces may have changed considerably, the ranking based on their binding energies is not significantly affected. Altogether, these results suggest that the relaxation protocol adopted in our previous study^13^ is sufficiently accurate in ranking the different binding modes by their binding energies.

**Figure 6:**
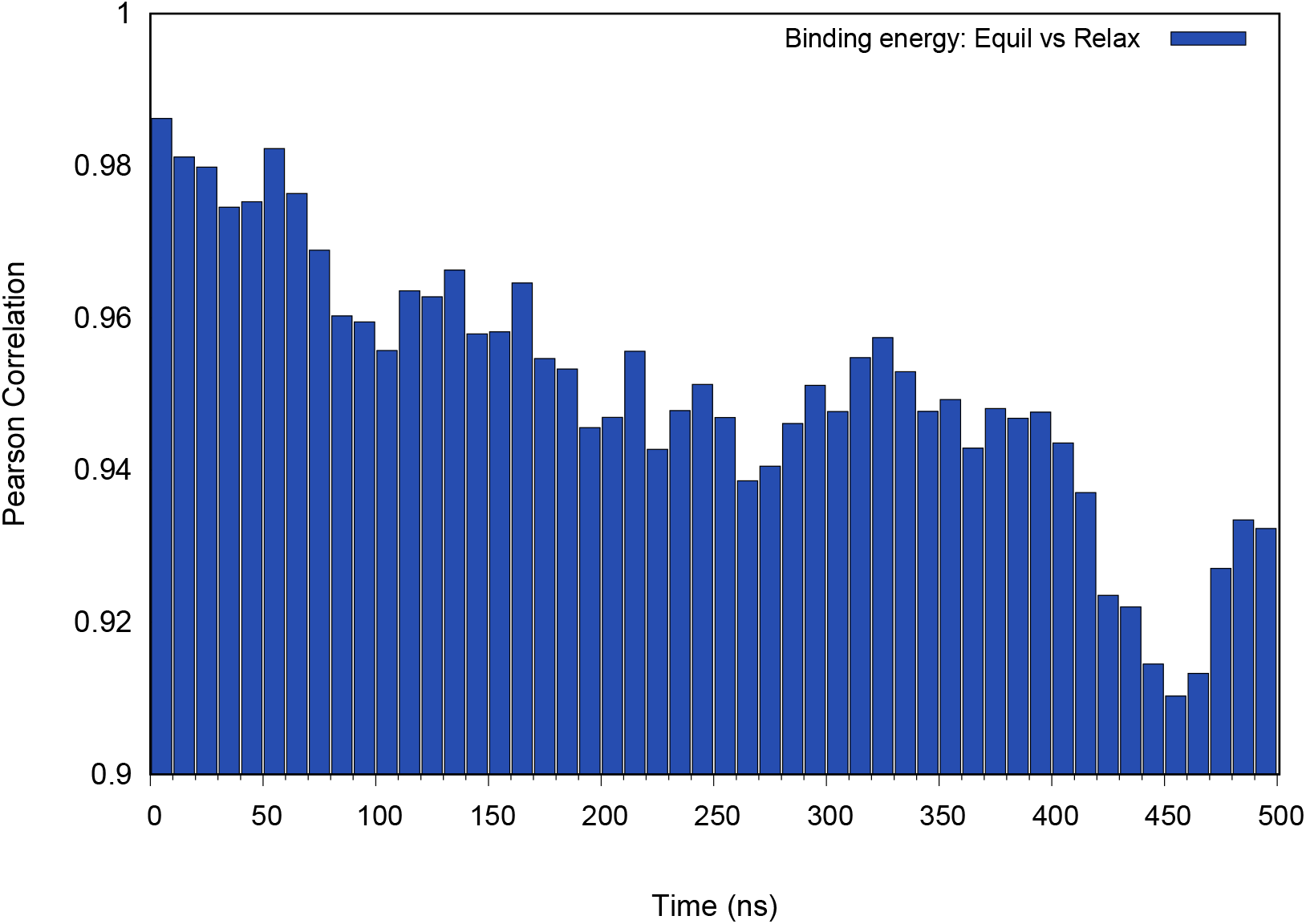
Pearson correlation coefficient between the average binding energy values of the long MD simulations and the short relaxation MD.^13^ The data of the 15 dimer set was used. The binding energies of the long MD equilibration replicates was averaged in intervals of 10 ns to show the correlation time evolution.

### Interfacial dynamics and polymerization modes

The results presented in the previous section show that the interfacial region of the considered dimers changed both in terms of energetic stability and residue composition during the 500 ns MD simulations. It is thus natural to ask if these modifications are able to drive different growth modes. In other words, does the growth mode change with time? In order to answer this question one needs to be able to determine the growth mode of any dimer conformation extracted from a MD simulation. By evaluating the polymerization angle (*θ*_*pol*_) and dihedral (*ϕ*_*pol*_) for each dimer conformation, we can then map them onto the growth landscape in the region (limited, unlimited or uncertain) that determines the growth mode (Figure 5). We recall that the latter was constructed by reducing monomers to spheres. For many conformations this approximation will work fine, but it breaks down when the dimer conformation deviates strongly from this geometric ideal, either due to rearrangements within the protein core or due to protrusions of the protein termini. In this case, visual inspection is needed to confidently classify the growth mode.

To characterize the interfacial dynamics and its relation with growth mode, we have averaged the polymerization angle *θ*_*pol*_ and polymerization dihedral *ϕ*_*pol*_ of the equilibrated segment of each replicate. We have then identified the dimer conformation within the ensemble of sampled dimers whose *θ*_*pol*_ and *ϕ*_*pol*_ more closely matches the average value. We term it the average conformation (or average dimer). With this approach, we can evaluate the dispersion of both angles during the simulation, and compare the results of the present simulations with those obtained in the shorter MD relaxation step reported in.^13^ Despite a few exceptions, there is an overall consistency in both simulations regarding the type of growth mode for the considered average dimers (Table 2 and Figure 7).

**Table 2.**
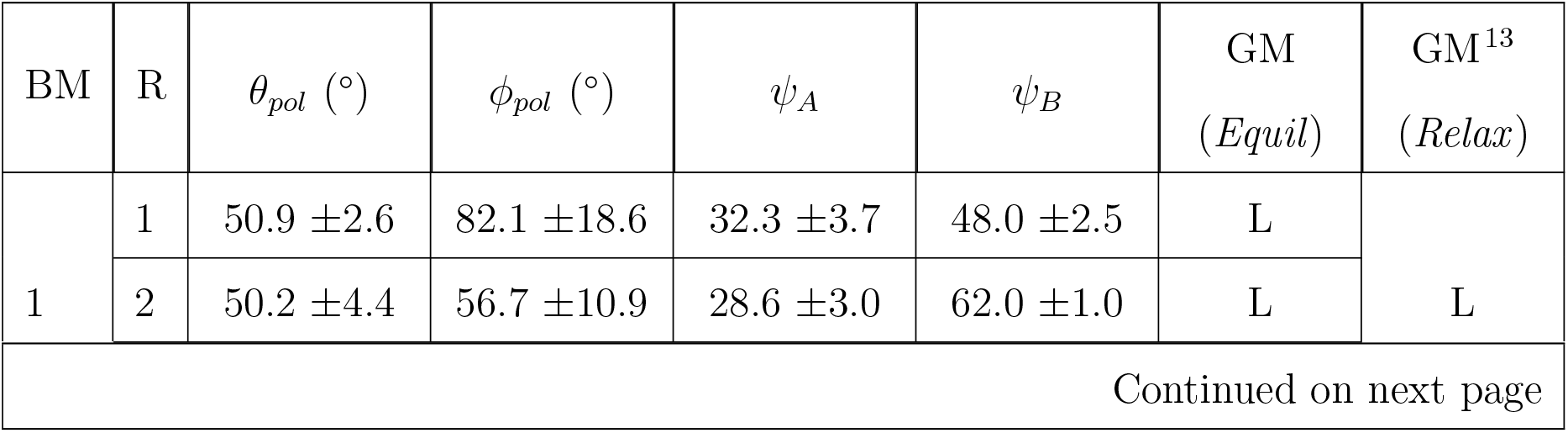
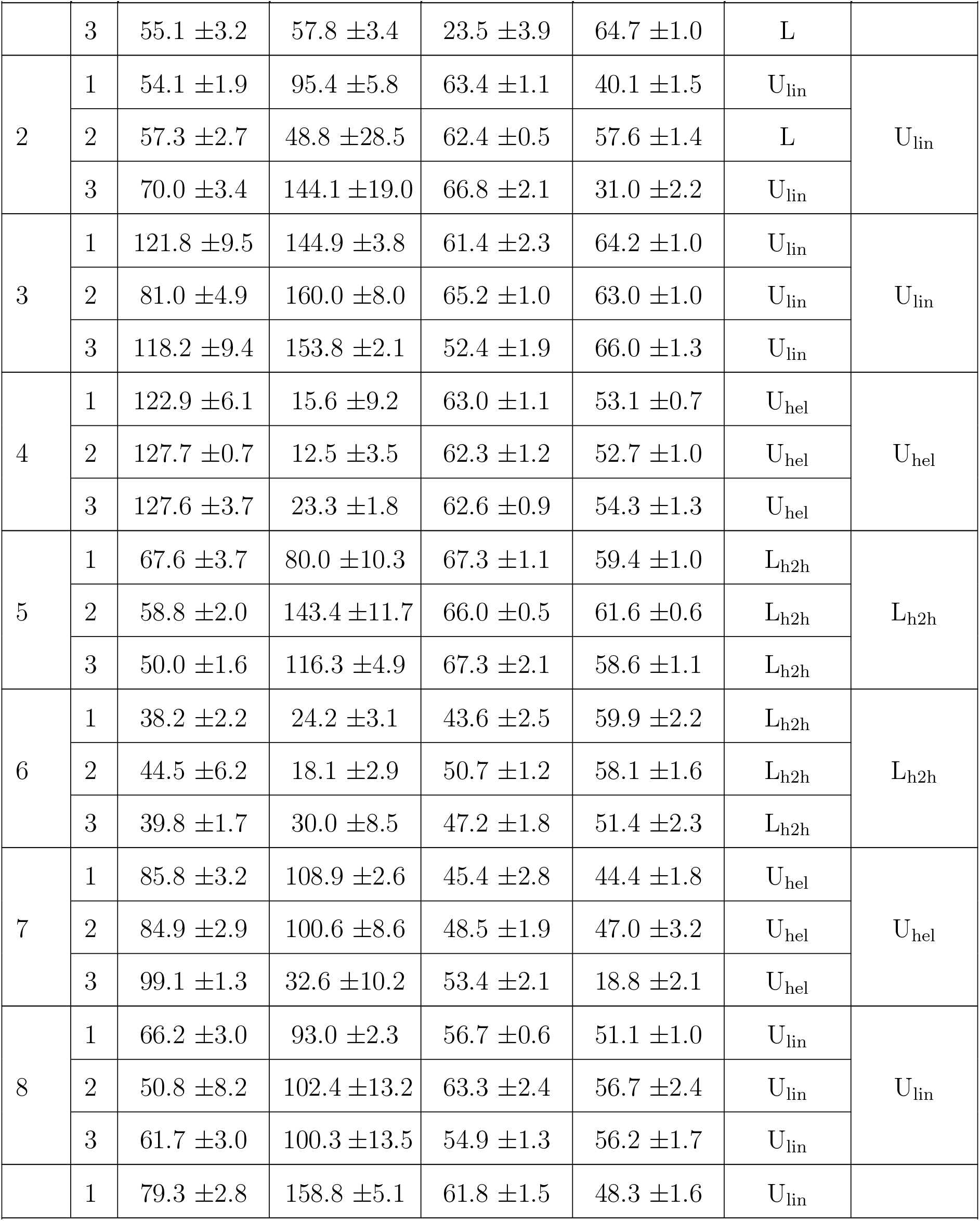
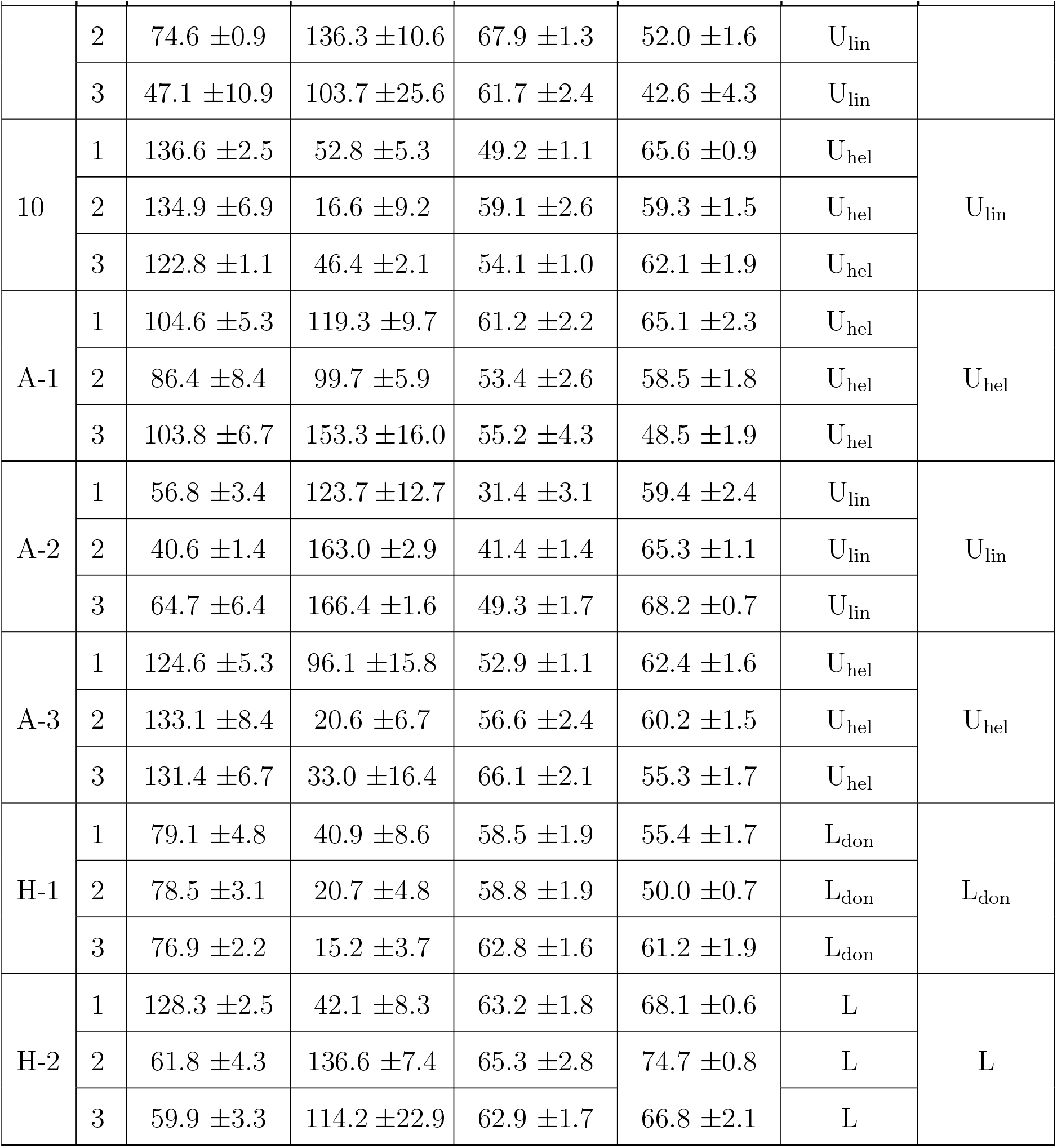
Average polymerization angle, *θ*_*pol*_, dihedral angle, *ϕ*_*pol*_, sphericity of monomers A (ψ_*A*_), and B (ψ_*B*_), and type of growth mode (GM) - Limited (L) or Unlimited (U), for the average structure (*Equil*). Within the L and U growth modes, we can further classify these structures as h2h (head-to-head), don (doughnut), hel (helix), and lin (linear) (see Figure 7 and Figure S6 of Supporting Information). Also shown is the GM (*Relax*) reported in our previous work.^13^ We show data for the 3 replicates (R1–R3) of the 10 most stable binding modes (BM1–10), the 3 binding modes with intermediate stability (A-1–A-3), and the 2 binding modes with lowest stability (H-1–H-2).

**Figure 7:**
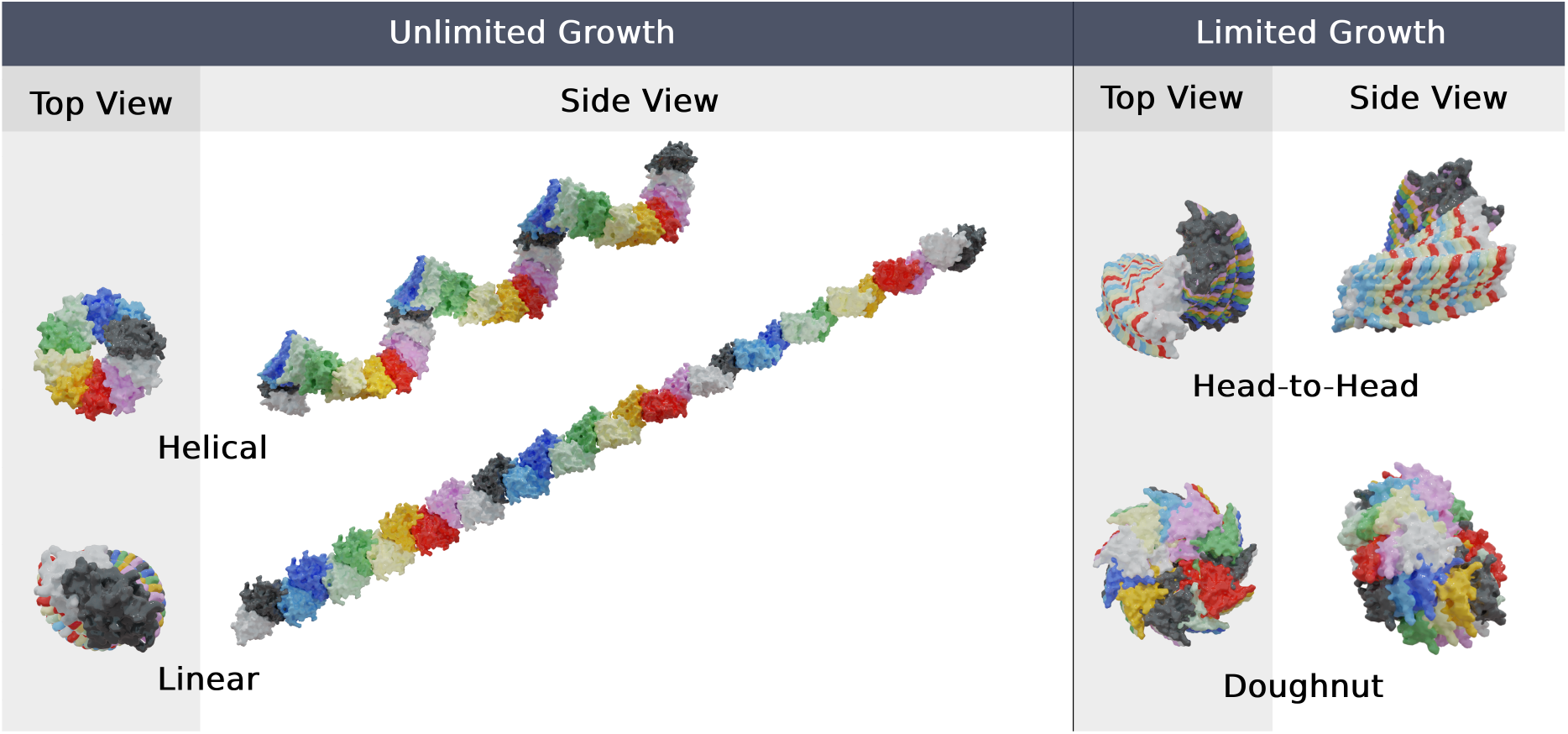
Examples of the different *β*_2_m dimer growth modes identified in our study. Each represented growth mode consists of 32 monomers. The BMs selected to illustrate the helical (U_hel_), linear (U_lin_), head-to-head (L_h2h_) and doughnut (L_don_) growth modes were BM-4 (R1), BM-3 (R1), BM-6 (R2), and H-1 (R2), respectively.

Our results show that, for the most stable binding modes within the analysed set (BMs 1– 10), the polymerization angle (*θ*_*pol*_) tends to change within a ∼30º range (Figure S5 of Supporting Information). This is expected since significant changes in the residue composition of the dimer’s interfacial region are usually required to trigger larger variations to the polymerization angle. An exception to this observation is illustrated by BM-3, which conserves only 61.2% of the original interface and shows a relatively high dispersion in the polymerization angles (large error values) for all replicates (Figure 8A). Nevertheless, the large *θ*_*pol*_ and *ϕ*_*pol*_ polymerization angles ensures a low number of clashes between monomers, leading to an unambiguous assignment of unlimited growth. A more detailed analysis of replicates 1 and 3 show a high *θ*_*pol*_ (∼ 120°) combined with a high *ϕ*_*pol*_ (∼ 150°), which lead to a linearized growth mode. Interestingly, the smaller *θ*_*pol*_ (∼ 80°) in replicate 2 caused a more pronounced rotation/distortion in the linear structure. These findings indicate that larger *θ*_*pol*_ angle values will favor less structured growth modes, eventually converging to fully linear and unlimited structures.

**Figure 8:**
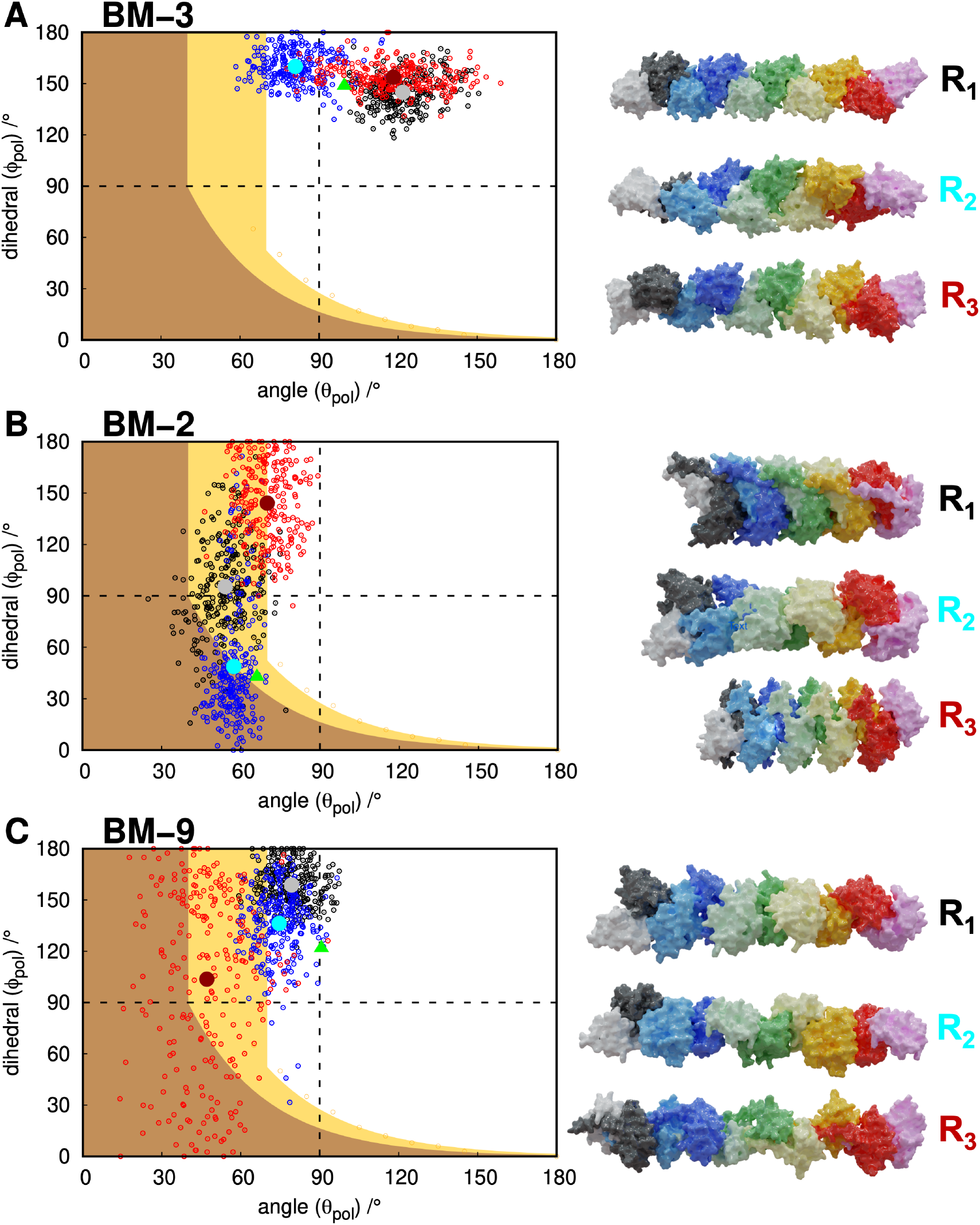
Growth landscape of BM-3 (A), 2 (B), and 9 (C). The black, blue and red dots represent the ensemble of 250 dimer conformations extracted from the equilibrated part of replicates 1, 2 and 3, respectively. The structure closer to the average *θ*_*pol*_ and *ϕ*_*pol*_ values of each replicate is shown in the corresponding color tone (grey, cyan and dark red). These are the structures used to generate the polymerization modes represented on the right hand side of each growth landscape. The green triangle represents the starting structure obtained from the initial MD relaxation step.^13^

The BM-2 is quite interesting owing to the structural sphericity of monomer B being significantly decreased (up to ∼ 30%) in replicate 1 and 3 (Table 2). Since this BM has a *θ*_*pol*_ ∼ 60°, it falls into the uncertain region of the polymerization landscape (Figure 8B). However, since monomer B, which is the one that is being consecutively added, has its detached termini in the dimers’ interfacial region, it deviates from the ideal spherical interface of the simple model. As a result, both replicates 1 and 3, have only a small structural overlap and are able to polymerize linearly, albeit being in a low *θ*_*pol*_ region. However, this sphericity deviation is smaller in replicate 2 and its limited growth prediction is quite clear.

The BM-9 also shows heterogeneity between replicates in their growth modes (Figure 8C). Similar to BM-2, we observe some loss of sphericity in monomer B, specially in replicate 3 (Table 2), due to the detachment of one of its termini that became part of the interfacial region. This leads to a decrease in the *θ*_*pol*_ polymerization angle, bringing some uncertainty to the growth mode viability. However, a visual inspection shows that, despite a few structural clashes that can be mitigated with small conformational rearrangements, this BM can easily grow in to an unlimited helical pattern.

## Conclusion

As we have observed in our previous work,^13^ *β*_2_m dimer structures are quite dynamic. Hence, it is reasonable to ask if structural modulations of the dimer’s interface (or binding mode), occurring in a timescale longer than 100 ns, can change the way they grow or polymerize. To explore this question, we developed a simple model that represents the geometry of the dimer’s interface by two angles, a polymerization angle (*θ*_*pol*_) and a polymerization dihedral (*ϕ*_*pol*_), combined it with the replication protocol, and calculated a (theoretical) growth landscape for idealized dimers (i.e., dimers formed by spherical monomers). The growth landscape is the space spanned by the two angles and contains three regions. A region corresponding to self-limited growth, a region corresponding to unlimited growth, and a region which we termed uncertain region. Given a dimer conformation sampled from a MD simulation trajectory, it suffices to determine its polymerization angles and map them on the growth landscape. When they fall into the uncertain region, an additional visual inspection step based on the replication protocol becomes necessary to establish the growth mode.

In the present study, we conducted long (3× 500 ns) MD simulations of 15 *β*_2_m dimer binding modes exhibiting different interfacial stabilities, starting from the relaxed conformations, which were previously obtained.^13^ With only two exceptions, all the considered dimers adopted more energetically stable interfaces, with stability increasing up to ∼23% for the top-10 dimers or ∼55% for the less stable ones. Despite these changes, the original ranking of BMs based on interfacial stability was not significantly altered. Since non-specific apolar interactions are the main drivers for these BMs,^13^ we continue to find a high correlation between the dimer stability and the interfacial area (*r* =0.93). However, one could not establish a clear relation between the percentage of conserved interfacial area (61-85%) and the corresponding change in interfacial stability (*r* =0.25), indicating that all these BMs are highly dynamic in nature, even the most stable ones.

To investigate if the growth mode changes within the time span of the MD simulations, an ensemble of 250 dimer conformations was extracted from the equilibrated part of each MD replicate (1 conformation/ns). For each conformation, we measured the polymerization angle and dihedral and mapped them onto the growth landscape. Since it is not feasible to replicate the dimer interface for the total 11250 dimer conformations, we took a conservative approach and considered the conformation whose polymerization angle and dihedral more closely match the averaged ensemble angles per replicate. This methodology also allows establishing some level of comparison with the results obtained in our previous study. ^13^ We noticed that for some binding modes (e.g. BM-9) the dispersion around the average is considerably large with different regions of the growth landscape being populated for the same replicate. This is a clear indication that the growth mode of the considered dimers can change within the ns–*μ*s timescale, contributing to the heterogeneity of the aggregation process. When considering the dimer conformation representing the ensemble average, we observe consistency with our previous results,^13^ which can be taken as an indication that a methodology that combines MC-ED with a standard relaxation MD protocol is able to deliver proper dimer models. Altogether, our approach led to a robust classification of which dimer configurations have limited and unlimited growth modes, with different consequences to their aggregation potential and biological impact.

## Author contributions

MM and PFNF designed research; NFBO and FEPR performed calculations; JNMV developed the MM/PBSA software and performed the calculations; NFBO, FEPR, and JNMV prepared the figures; all authors analyzed the data; NFBO, FEPR and JNMV prepared a preliminary version of the manuscript; PFNF and MM wrote final version.

## Data and Software Availability

The GROMACS package is freely available software used to perform MD simulations and can be downloaded at https://manual.gromacs.org/documentation/2018.6/download.html. PyMOL v2.5 is also free software for molecular visualization and generating high quality images. It can be downloaded from https://pymol.org/2. Blender is a free and open-source 3D creation suite, licensed as GNU GPL, and it can be downloaded from https://www.blender.org/

## Acknowledgements

We acknowledge financial support from Fundação para a Ciência e a Tecnologia through grants CEECIND/02300/2017, 2021.06409.BD, 2021.05909.BD, and 2022.11124.BD, and projects PTDC/FIS-OUT/28210/2017, UIDB/04046/2020, and UIDP/04046/2020.

## Supporting Information Available

RMSD, interface area, secondary structure, and binding energies for *β*_2_m dimers. Polymerization landscape of all dimer configurations. Growth modes graphical representation for all dimers.

## TOC Graphic

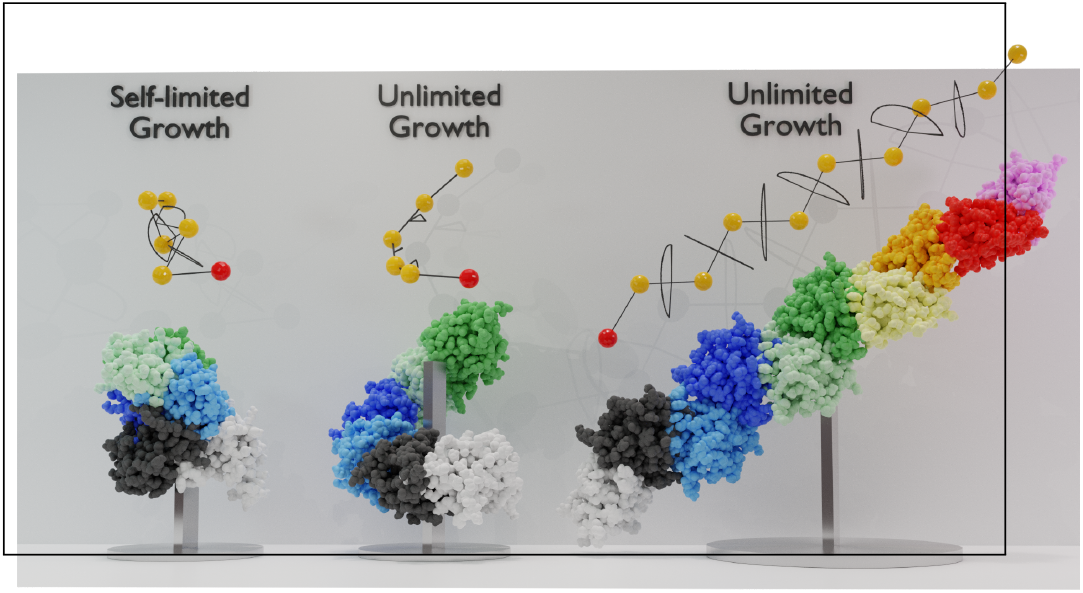

