## Supporting information for "Interfacial dynamics and growth modes of *β*_2_-microglobulin dimers"

### Supporting Information: Interfacial dynamics and growth modes of $\beta_2$-microglobulin dimers

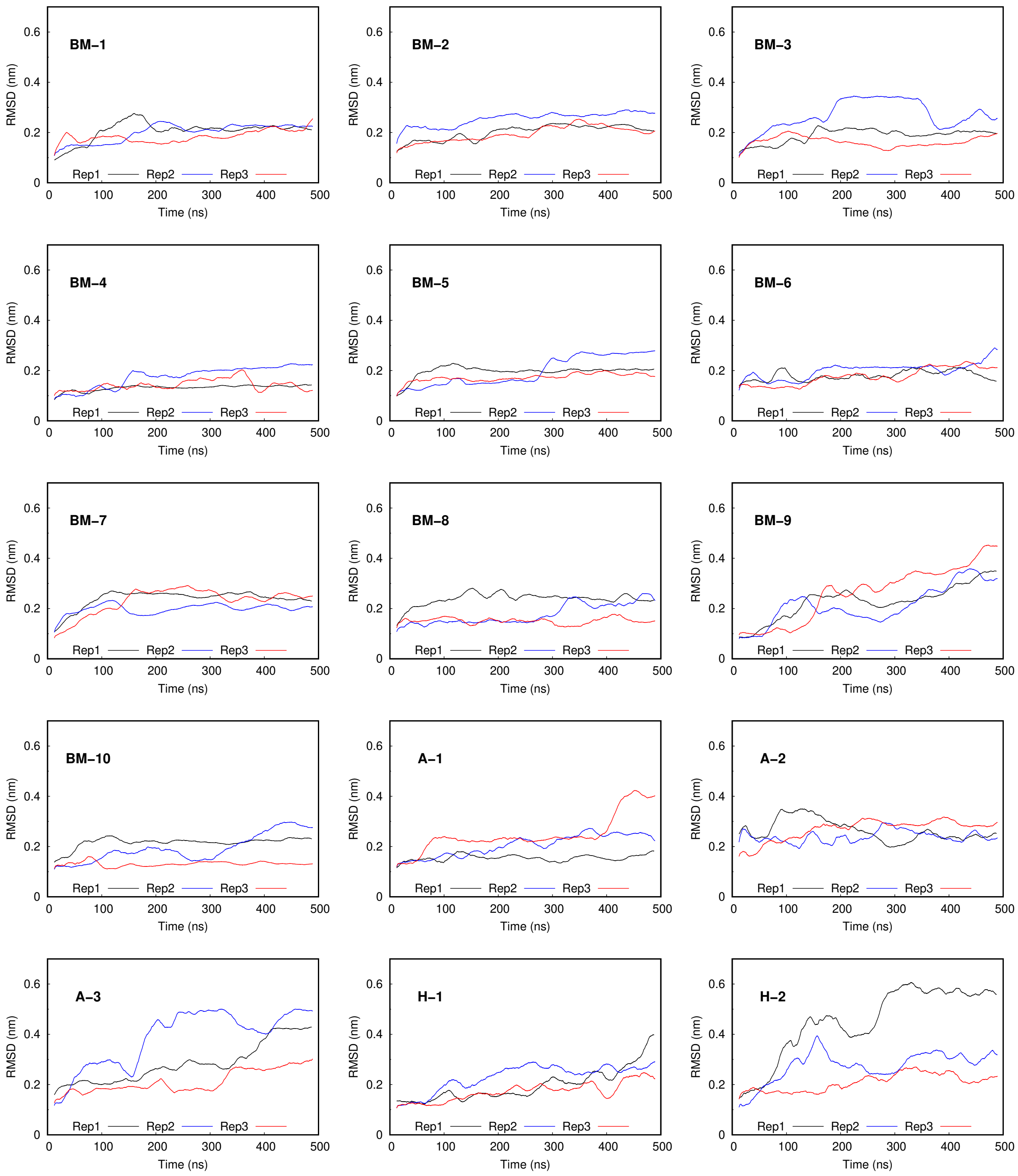


Figure S1: Dimer RMSD values of the 3 replicates of each dimer configuration studied.

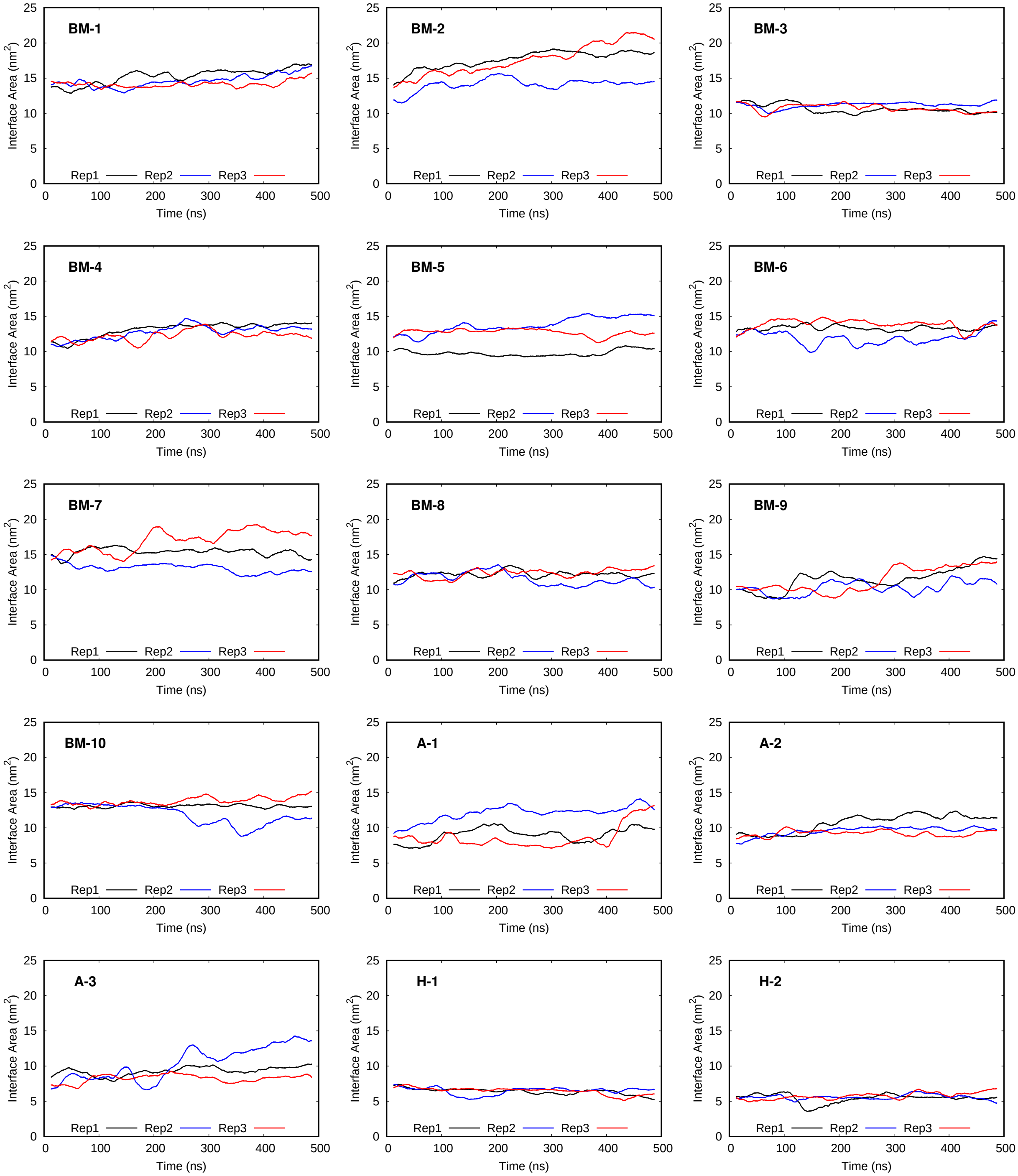


Figure S2: The evolution of interfacial area for the 3 replicates of each studied binding mode.

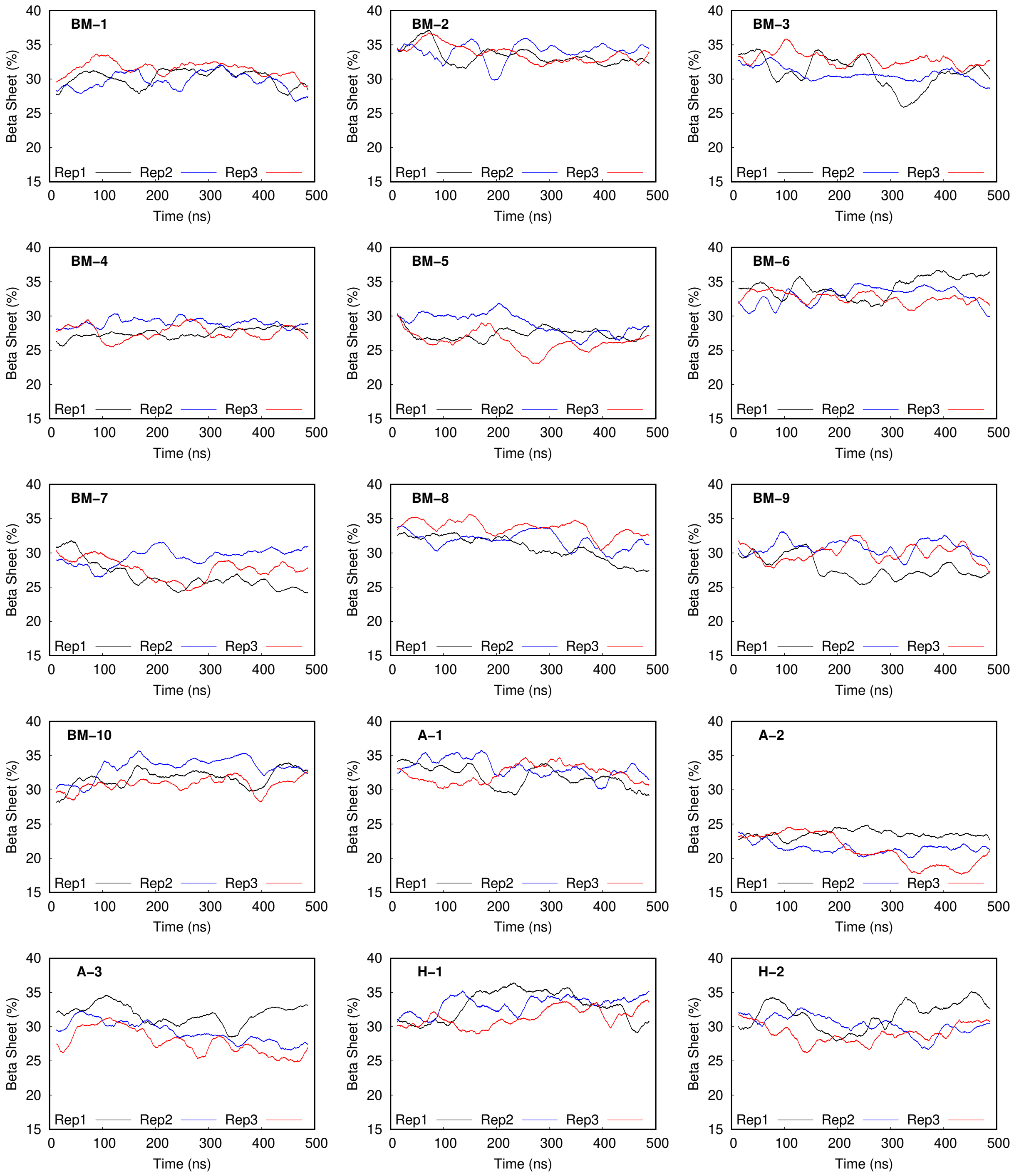


Figure S3: The evolution of beta-sheet percentage for the 3 replicates of each studied binding mode.

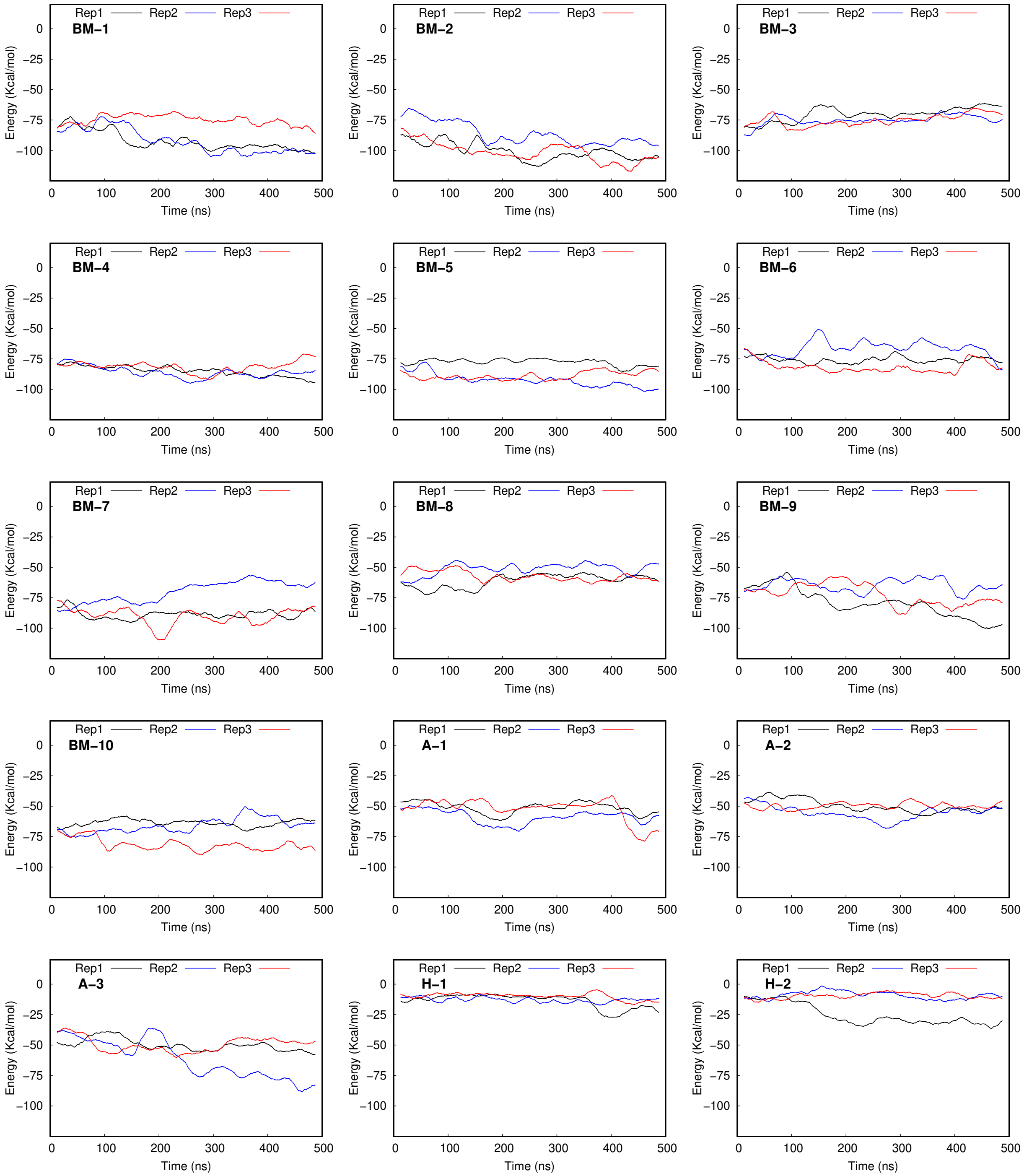


Figure S4: Time evolution of binding energy for the 3 replicates of each studied binding mode.

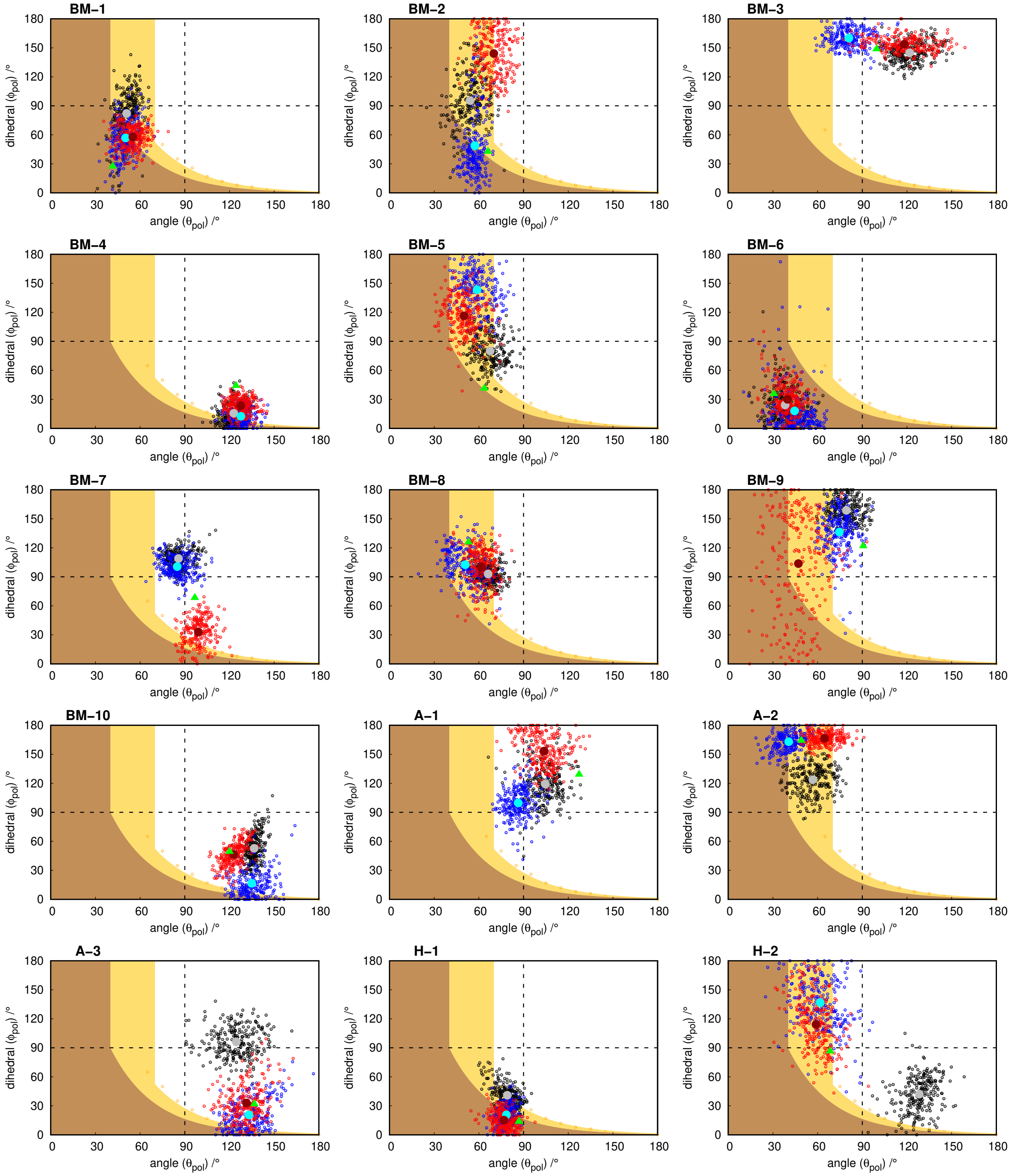


Figure S5: Growth landscape as a function of $\theta_{pol}$ angle and $\phi_{pol}$ dihedral angle. The black dots represent the upper limit of angle/dihedral combinations that yield a limited growth polymer as predicted by the simple model. The orange region represents the region where polymer growth is limited, the yellow color represents the uncertain region (one that requires visual inspection), and the region in white represents the unlimited growth.

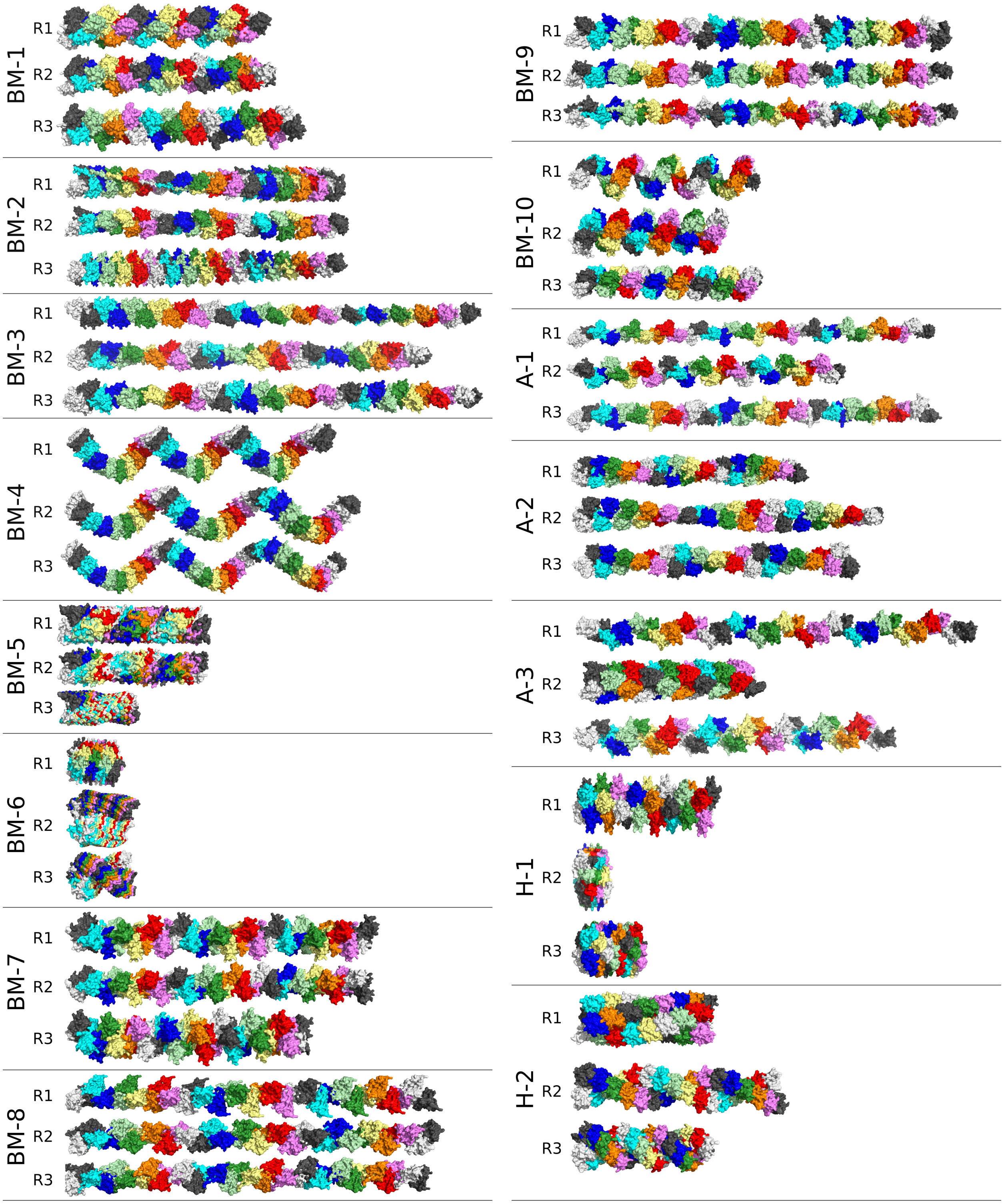


Figure S6: Polymerization growth mode side-view representation for all the studied binding modes. Each polymer consists of 32 subunits (in repeating different colors).
